# Genomes of Animal Mitochondria have not Evolved According to Common Models of Substitution Probabilities

**DOI:** 10.1101/2020.04.26.062596

**Authors:** James Cavender

## Abstract

Common probabilistic models of substitutions of bases (Jukes-Cantor, Kimura 2-parameter, Tamura-Nei, F84, HKY, and the 6-parameter models used in linear invariants methods) must be rejected, at least for mitochondrial genomes of animals. They are rejected by a new test that is simple and lenient.

## 1. Introduction

It would be impossible to infer any aspect of an evolutionary tree from nucleic acid sequences without some model of the evolution of sequences. Here I treat the Jukes-Cantor model (Felsenstein 2004, p. 156), the Kimura two parameter model (Felsenstein 2004, p. 196), the Tamura-Nei model, F84, and HKY (Felsenstein 2004, p. 200 *et seq*.). These are all special cases of models used in linear invariants methods. For mitochondrial genomes of animals, I will reject the less general models by rejecting linear invariants models.

People who derive methods of phylogenetic inference from modeling assumptions are showing that those assumptions are sufficient. They sometimes don’t investigate whether their assumptions are necessary. Here I show that if any of the models just mentioned is necessary for a method to succeed, then that method will not succeed, at least for mitochrondrial genomes of animals.

The method of linear invariants for inferring evolutionary trees from nucleic acid sequences has the advantage that it depends for its success on very weak assumptions about the way the sequences evolve. The mathematical theory is presented in Lake 1987, Cavender 1989, 1991, and Nguyen and Speed 1992. (But Lemma 2 of Cavender 1991 is wrong.)

## 2. Necessary Conditions of Linear Invariants

In this paper, I treat two-species trees with a root species `R` and leaf species `S` and `T`. For a particular site in a nucleic acid sequence, the Markov matrix,

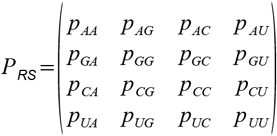

says that the conditional probability of finding G in `S` given A in the root `R` is *p_AG_* and similarly for the other components of the matrix. (I am using the convention that U means “U or T, depending on your acid”). A matrix *P*_RT_ is defined analogously. The linear invariants method requires these matrices to have the form

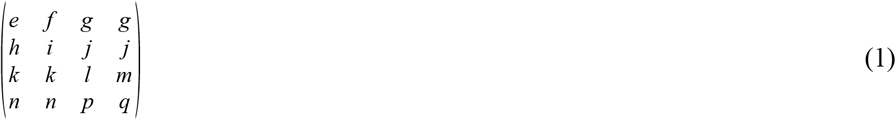

and satisfy the two constraints,

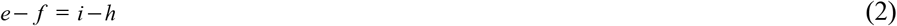

and

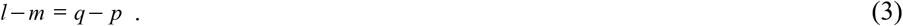

Each site in the sequence may have its own Markov matrices *P*_RS_ and *P*_RT_ satisfying (1), (2), and (3). The probabilities of A, G, C, and U in the root may be anything. Different sites are assumed to be statistically independent. Yes, (1), (2), and (3) look strange, arbitrary, and implausible at first. But the popular Kimura two-parameter model satisfies (1), (2) and (3), so if you can accept that model you may not reject this much less restrictive one. And mathematically (1), (2) and (3) are not arbitrary. They specify a maximal semigroup of Markov matrices under which linear invariants will work, that is, necessary conditions. This the *balanced transversion* model.

There is a family of alternatives, the *nonbalanced transversion* models (or “beyond balanced transversion” models). In these, (1), (2) and (3) are replaced by

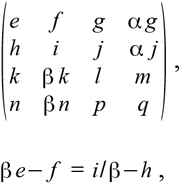

and

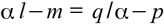

where α and β are positive constants. From Felsenstein 2004 (equation 13.11, P. 203) one can deduce that the Tamura-Nei model, F84, and HKY are nonbalanced models.

## 3. Invariants

Two aligned sequences determine a *spectrum*,

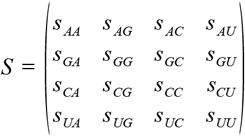

in which *s_AG_* is the number of sites in the alignment where an A in the first sequence aligns with a G in the second, and similarly for the other components. With (1), (2), and (3) in force, the forms

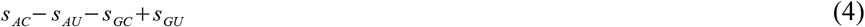

and

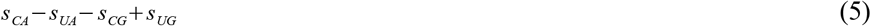

are random variables with mean zero. In more elaborate language, let

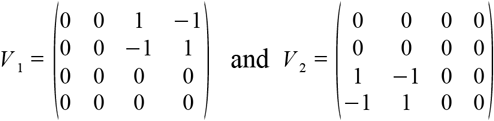

and define an inner product by

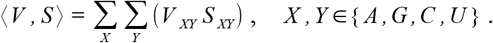

Then (4) is 〈*V*_1_, *S*〉, (5) is 〈*V*_2_, *S*〉, and *V*_1_ and *V*_2_ are two *linear invariants* that form a basis of the two-dimensional vector space of all linear invariants of the two-species tree. The nonbalanced transversion models also have linear invariants.

## 4. Results

I studied mitochondrial genomes because these appear to be long enough to provide statistical significance.

Table 1 shows the spectrum for the complete mitochondrial genomes of two species of Cnidaria. Species `S` and `T` are the brown hydra *Hydra oligactis* (GenBank Accession NC_010214) and the brown sea anemone *Metridium senile* (NC_000933.1). I copied the sequences from GenBank and used the Smith-Waterman algorithm of EMBOSS (https://www.ebi.ac.uk/Tools/psa/emboss_water/nucleotide.html) to align them. The entry 275, for example, means that at 275 sites, an A in the *Hydra* sequence aligned with a C in *Metridium*. For this spectrum, (4) is −54, not zero. If (4) and (5) are significantly far from zero, I must reject (1), (2), and (3) as a hypothesis.

**Table 1.** A spectrum

|  |  |  |  |  | First<br>sequence |
| --- | --- | --- | --- | --- | --- |
|  | AA | AG | AC | AU | A |
|  | 2899 | 439 | 275 | 355 | 3968 |
|  | 25.33% | 3.84 | 2.40 | 3.10 | 34.67 |
|  | GA | GG | GC | GU | G |
|  | 84 | 1066 | 65 | 91 | 1306 |
|  | 0.73 | 9.31 | 0.57 | 0.80 | 11.41 |
|  | CA | CG | CC | CU | C |
|  | 72 | 83 | 859 | 103 | 1117 |
|  | 0.63 | 0.73 | 7.51 | 0.90 | 9.76 |
|  | UA | UG | UC | UU | U |
|  | 317 | 340 | 343 | 4054 | 5054 |
|  | 2.77 | 2.97 | 3.00 | 35.42 | 44.16 |
| Second sequence: | 3372 | 1928 | 1542 | 4603 | 11445 |
|  | 29.46 | 16.85 | 13.47 | 40.22 |  |

In more correct terms, I want to test whether the *Hydra-Metridium* spectrum lies significantly far from the subspace that is orthogonal to *V*_1_ and *V*_2_. To do this, set *Y*=*s_AC_*+*s_GU_* and *n_Y_*=*s_AC_*+*s_AU_*+*s_GU_* so that under the null hypothesis *Y* is binomially distributed with parameters *n_Y_* and ½. Set

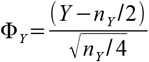

so that Φ_*Y*_ is approximately standard normal. Compute *Z* and **Φ**_*Z*_ from *s_CA_*+*s_UG_* and *s_CA_*+*s_UA_*+*s_CG_*+*s_UG_* similarly. Set 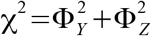 to form a chi-square test statistic with two degrees of freedom. In this case, χ^2^=4.505 which is not large enough for rejecting the null hypothesis. (The rejection levels for a chi-square test with 2 degrees of freedom are 5.99 and 9.21 at the 5% and 1% levels, respectively.) Strictly, the modeling assumption that each base in the sequence has its own (1), (2), and (3) is not compatible with this statistical analysis. The distributions of *Y* and *Z* are mixtures of binomials rather than binomial. As a practical matter, this means the power of the test is reduced but the size is not impaired. That is, acceptances should not be trusted but rejections can be. (But this is ordinarily the case with hypothesis tests.)

Repeating this analysis with the freshwater jellyfish *Craspedacusta sowerbyi* (JN593332) and the jellyfish *Rhopilema esculentum* (NC_035740) gives χ^2^=23.812. This is decisive grounds for rejection. But this spectrum exactly satisfies the invariants of the nonbalanced transversion model with α=3.83 and β=0.188 and also for the model with α=0.6642 and β= 1.1156. Two nonbalanced models like these can be found for most spectra, as follows.

From Cavender 1991, page 73, or Cavender 1989, page 313, these matrices are a basis of the invariant space under the nonbalanced model:

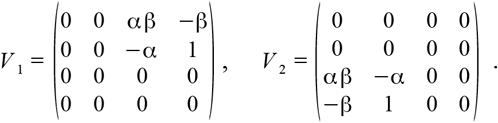

Thus we seek α and β that, for the given *s*, satisfy

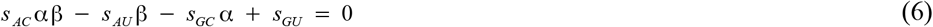

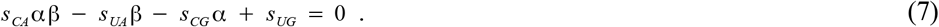

this system is easy to solve if *s_AC_* or *s_CA_* is 0. Otherwise, eliminate αβ by adding equations. That is, add

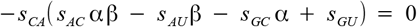

to

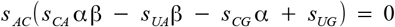

obtaining

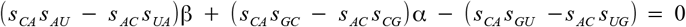

or *h*β + *k* α = *j* with *h, k*, and *j* defined in the obvious way. Then

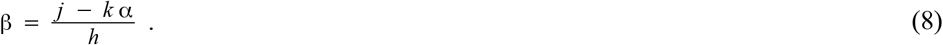

Substituting (8) into (6) gives

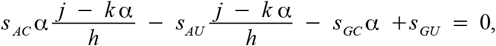

a quadratic equation to solve for α. Finally, recover β using (8). But the quadratic equation that gives two solutions might not always give real, positive α and β, if only because spectra are random objects.

The chi-square computation just given for the balanced transversion model can be modified for use with nonbalanced transversion models. Is there a nonbalanced transversion model under which neither of these Cnidarian spectra is rejected by the chi-square test? No. The best is surprisingly close to balanced, with α=0.842 and β=0.963 but has χ^2^=37.94 for both spectra, requiring rejection at the 1% level. Fudging parameters to minimize a χ^2^ statistic is another dubious procedure. Again, it reduces power but does not impair size.

This analysis of Cnidaria can be repeated for any taxon containing four species whose mitochondrial genomes have been sequenced. I have done it for 11 phyla, proposed phyla and former phyla. The results are shown in Table 2. The balanced model is rejected for almost every phylum. I hoped to find an α and β that would work throughout Metazoa, but they don’t exist. No consistent α and β to account for evolution in the various phyla showed up. No linear invariants model is correct. (For Ctenophora no credible nonbalanced model fit both spectra. The minimum-seeking algorithm halted while pursuing α and β to ridiculous extremes. This is doubtless because the mtDNA of *Mnemiopsis leidyi* is anomalous, as Pett, *et al*. 2011 discovered.)

**Table 2:**
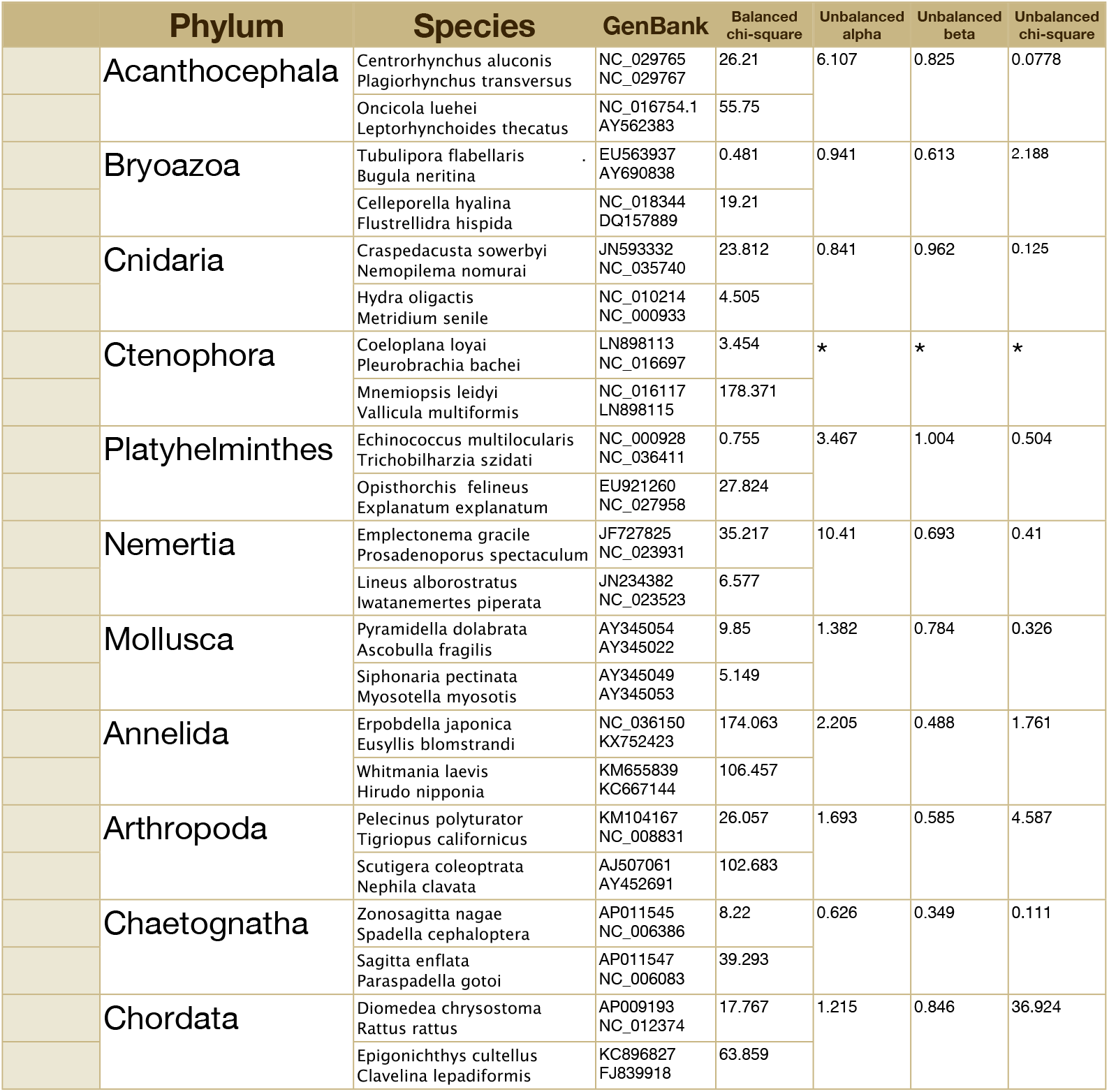
Fit of models to spectra

## 5. Discussion

The failure of mtDNA to comply with linear invariants models is not because different parts of the DNA evolve at different rates and it is not because different rates apply on different edges of the tree. The linear invariants models allow that. (The tests in Goldman 1993 do not.) The rejections happened because (1), (2), and (3) are false hypotheses, or possibly because of dependence among substitutions at different sites.

One might object that it is not plausible that the balanced model is *exactly* right, and therefore any sufficiently powerful test is bound to reject it. But my test is hardly powerful! Moreover, Table 2 shows that perfect balance is more than a little false, according to my best estimates.

The models that I have rejected, all proposed before 1992, look like very sound applied probability. It is surprising to find that molecular evolution is so non-uniform, especially in mitochondria, so uniform in their environments and functions. These 22 spectra do not tell us whether their divergence from reasonable models indicates grand, stately flows, or rapid, wide fluctuations. With a careful choice of mitochondrial genomes, it should be possible to tell. Many of the investigators who produced the 44 sequences that I have studied here did so in order to clarify phylogenetic relations. This might be a futile program.

Mitochondrial genomes of animals are CG-poor, as seen in Table 1, for example. To maintain this imbalance over time requires small α and large β, if in each edge of the tree all sites in the sequences are governed by the same Markov matrix. Since this is not seen, I conclude that a large majority of the Metazoan mitochondrial sequence is CG-poor and very stable. The minority is most of what I have analyzed here. This is a new line of evidence for old knowledge (Aquadro, Kaplan and Risko 1984).

